# The quorum sensing system NprR-NprRB contributes to spreading and fitness in colony biofilms of *Bacillus thuringiensis*

**DOI:** 10.1101/2021.01.24.428019

**Authors:** Abel Verdugo-Fuentes, Mayra de la Torre, Jorge Rocha

## Abstract

Quorum sensing (QS) are intercellular communication mechanisms to coordinate bacterial gene expression in response to signaling molecules. In *Bacillus thuringiensis* the QS system NprR-NprRB (receptor protein-signaling peptide) regulates the expression of genes related to nutrient scavenging during necrotrophism and also modulates sporulation onset. However, the relevance of QS in free-living stages of *B. thuringiensis* is less known. In this work, we depict the contribution of this QS system to spreading in colony biofilms. Through a spreading assay in spotted colonies of *B. thuringiensis* Bt8741 Wt and derived mutants, we find that the spreading phenotype depends on the NprR regulator and on the extracellular signaling NprRB peptide. We also show that this phenotype is associated to an increased fitness of the bacterium in these experimental conditions. Exogenous addition of a lipopeptide surfactant was sufficient to recover spreading in the Δ*nprR-nprRB* mutant, indicating that the phenotype could be mediated by the lipopeptide kurstakin. Finally, we suggest that the spreading is relevant in nature, since it occurs in the sole presence of soil nutrients, and it is conserved in several species of *Bacillus* commonly found in soil. This novel function of NprR-NprRB highlights the relevance of this QS system on the evolution and on the free-lifestyle ecology of *B. thuringiensis*.

## 1. Introduction

Bacteria from the genus *Bacillus* are widely distributed in aquatic and terrestrial ecosystems. Some species are relevant for the study of cell differentiation since they form spores to survive in hostile environments (1, 2). Other species are also economically relevant, and many strains have been used for producing biotechnological goods such as lipopeptide biosurfactants (3), endotoxins (4, 5), hydrolytic enzymes (6), bacteriocins (7), flavors (8) and probiotics (9). A notorious example is *Bacillus thuringiensis* (10, 11), which produces crystalline inclusions comprising Cry and Cyt proteins that are toxic for a wide range of insect orders. For this reason, *B. thuringiensis* has been used for biological control of agricultural pest and for induction of plant growth, for more than 50 years (5, 12, 13).

*B. thuringiensis* was initially isolated from dead insects and from soil, but it has also been found in phyllosphere and as endophyte of plants (14, 15). However, research efforts are mostly focused on the saprophytic lifestyle of *B. thuringiensis* in insects or on *in vitro* cultivation in shaken flasks or reactors for biological control of pests (16–18). In fact, *B. thuringiensis* and other species from the *Bacillus cereus* group have been widely used as model for the study of the intricate control of virulence, necrotrophism and sporulation (10, 19–21). These behaviors are modulated by quorum sensing (QS), which allows individuals in a population to perform collective functions, by sensing signal molecules synthesized and secreted by other bacterial cells (22, 23). In the *B. cereus* group, the RRNPP family of QS receptors (Rgg, Rap, NprR, PlcR, and PrgX), are activated by small, cognate signaling peptides encoded in their genomes (24, 25). These receptor-peptide QS systems carry out diverse functions, e.g. PlcR acts as pleiotropic regulator of virulence (26), while NprR and the repertoire of Rap proteins are multifunctional regulators that interconnect several differentiation programs such as sporulation, nutrient scavenging and biofilm formation (27–29).

Nevertheless, little is known about *B. thuringiensis* ecology in free lifestyle (30–32) or its effect on the microbial population and communities on the phyllosphere or as endophyte (13, 33, 34). *B. thuringiensis* could alter the microbial populations in nature due competition for nutrients and space. Likewise, the toxic crystal proteins produced by this bacteria could be used as substrate by other organism in the same habitat, or may be toxic to others (35). Accordingly, the complex regulatory circuits that direct differentiation in dead insects (or in culture media) may play relevant roles in other natural settings. Indeed, these regulatory QS systems are essential for individuals to cooperate and form multicellular structures like biofilms and fruiting bodies, to jointly scavenge for resources, to attack competitors, for predatory group behaviors, to defend themselves against predators and harsh environments, or to move together across surfaces (36–38).

In this work, we find that the QS system NprR-NprRB controls spreading in colony biofilms of *B. thuringiensis* Bt8741 in agar media. For this, we developed an experimental system using spot-inoculated strains of wild-type and derive mutants of Bt8741 strain on agar media. We find that the spreading phenotype is regulated by NprR and requires specific signaling by the NprRB heptapeptide. We also show that spreading allows increased growth and fitness in colony biofilms of Bt8741; furthermore, our results suggest that this new function is mediated by the production of a lipopeptide surfactant. Finally, we speculate that spreading could occur in nature, which highlights the relevance of the regulatory systems that coordinate this collective trait, in the evolution and ecology of *B. thuringiensis* and other *Bacillus* species.

## 2. Results

### 2.1 Spreading is robust to different experimental conditions in B. thuringiensis

We found a spreading phenotype in colonies of *Bacillus thuringiensis* strain Bt8741, that could be associated to motility of *B. thuringiensis* cells in nature, and to colonization of insect or plant hosts (30). In order to contribute to the knowledge of the free-living ecology of *B. thuringiensis*, spreading of strain Bt8741 Wt was assessed under different agar concentrations, nutrient dilutions and culture media. The strain spread at all agar concentrations tested (Figure 1a), but the morphology and size of colonies varied, being the highest at 1.5% agar. Bt8741 strain spreading decreased at higher dilutions of nutrient broth (Figure 1b); therefore, growth may be is important for this phenotype. This behavior was also consistent in different culture media (Figure 1c). Bt8741 colonies exhibited the maximum spreading in BHI, which contains the highest concentration of organic nitrogen (proteose peptone 10g/L, plus brain and heart infusion) among the tested media. It was followed by TSA (soy peptone 20g/L), LB (casein peptone 10g/L) and nutrient broth (beef peptone 5 g/L). These results show that spreading of Bt8741 colonies is robust to variable environments and were useful to define the experimental conditions in the following stages of this work.

**Figure 1.**
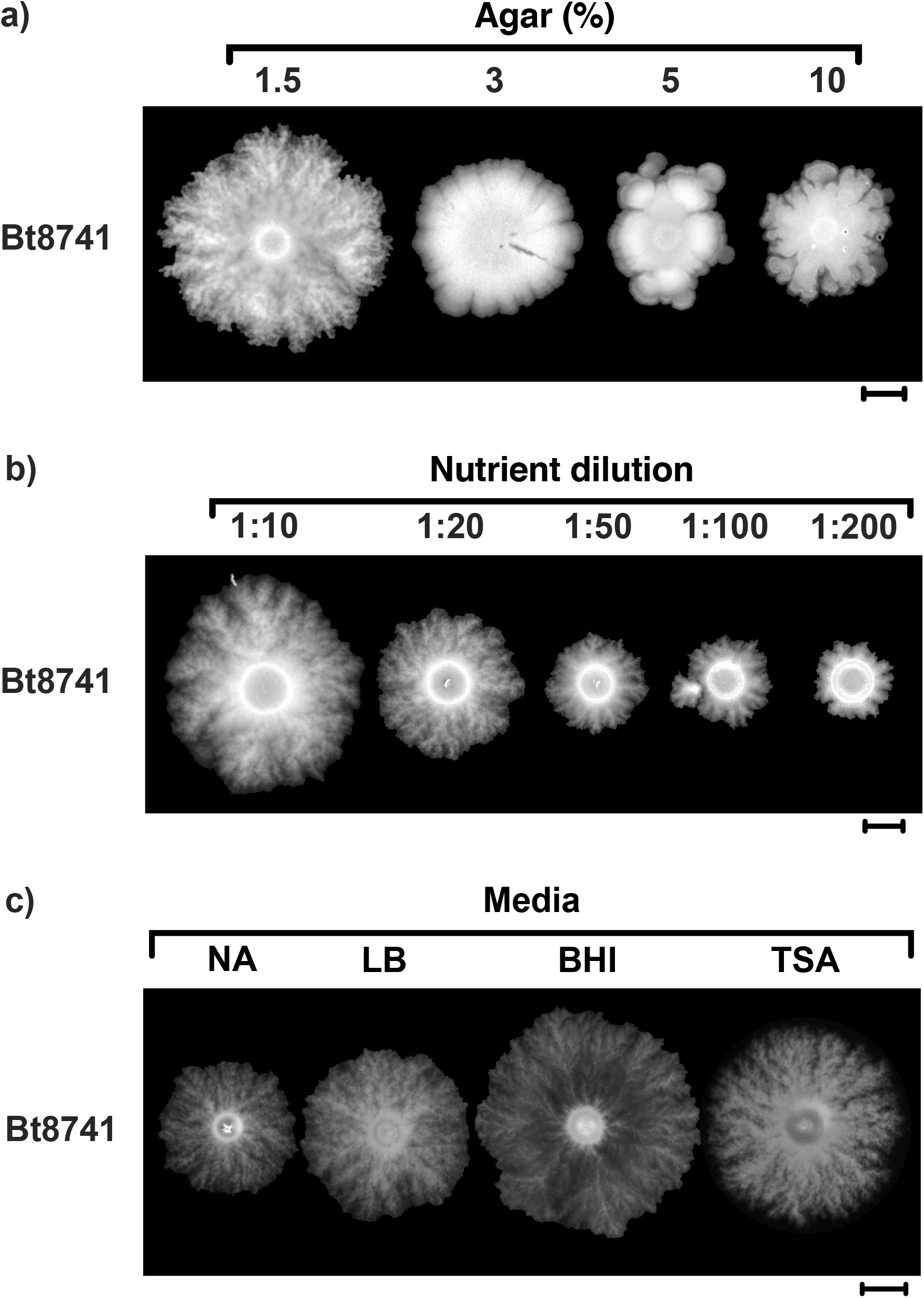
Spreading is robust to different experimental conditions in *Bacillus thuringiensis* Bt8741. (**a**) The spreading phenotype occurs in different agar concentrations (using nutrient broth). (**b**) The spreading phenotype varies with nutrient dilution (using 1.5 % agar; 1X nutrient agar contains 8 g/L of nutrient broth). (**c**) Spreading of Bt8741 in different culture media at 1.5% agar. NA, nutrient agar; LB, Luria Bertani agar; BHI, brain hearth infusion agar; TSA, tryptic soy agar. The scale bar indicates 5 mm.

### 2.2 Spreading in Bt8741 is associated to the NprR-NprRB QS system

In previous work, we have depicted the functions of NprR-NprRB and other quorum sensing (QS) systems using the laboratory strain Bt8741 (28, 39, 40, 43). During rutinary culturing in agar plates, we noticed that a one week-old colony of the the *ΔnprR-nprRB* mutant was defective in spreading; therefore, we decided to investigate the participation of NprR-NprRB in this phenotype. We followed the spreading of spot-inoculated colonies of Bt8741 Wt strain, Δ*nprR-nprRB* mutant, Δ*nprR-nprRB* (pMAD-NprR) and Δ*nprR-nprRB* (pMAD-NprR-NprRB) complemented strains (40) (Supplemental Table S1), for 7 days in nutrient agar plates. After 24h, we observed that all strains grew similarly. However, at days 3-7, we confirmed that spreading of the *ΔnprR-nprRB* mutant was severely decreased compared to the Wt strain (Figure 2a). Interestingly, the Δ*nprR-nprRB*(pMAD-NprR-NprRB) complemented mutant recovered the spreading phenotype, but the Δ*nprR-nprRB*(pMAD-NprR) strain –that carries the NprR receptor but lacks the gene coding for the signaling peptide –did not (Figure 2a). Using a zoom microscope, we imaged the edge of spreading and non-spreading colonies from this experiment. We found that spreading was related to the presence of ‘dendritic’ projections, probably composed of chained or elongated cells, in the Wt and the Δ*nprR-nprRB*(pMAD-NprR-NprRB) complemented strain. These projections were absent in the non-spreading Δ*nprR-nprRB* mutant and Δ*nprR-nprRB*(pMAD-NprR) complemented strain. This experiment show that the NprR-NprRB QS system is involved in the control of spreading in Bt8741.

**Figure 2.**
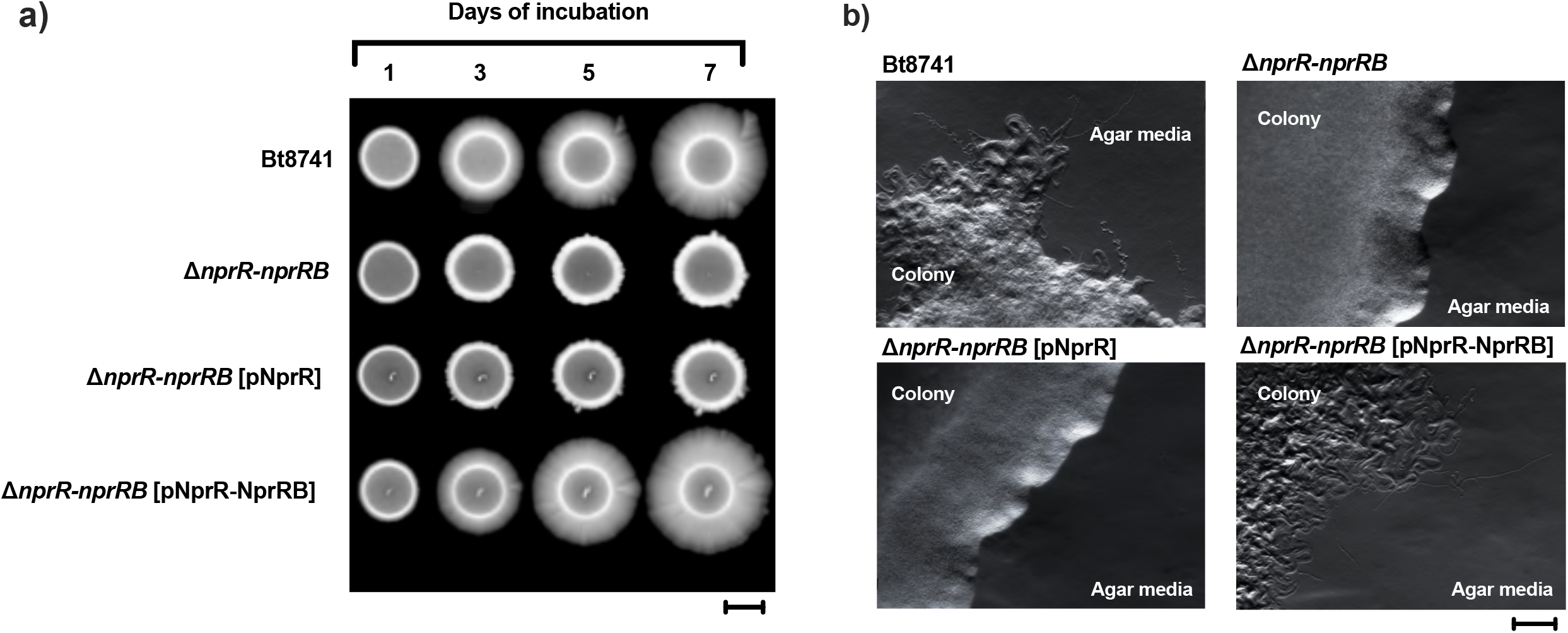
Spreading phenotype in *B. thuringiensis* Bt8741 is associated with the NprR-NprRB QS system. (**a**) Spreading in colony biofilms of Bt8741 Wt, Δ*nprR-nprRB* mutant, Δ*nprR-nprRB* (pMAD-NprR) and *B. thuringiensis* Δ*nprR-nprRB* (pMAD-NprR-NprRB) complemented strains was individually assessed. Plates were imaged at different days using a gel documentation system. Scale bar, 5 mm. (**b**) Edge of colonies from spreading assays. Spreading colony biofilms were imaged using a stereo microscope. Scale bar, 0.2 mm.

### 2.3 Extracellular signaling by NprRB mature peptide is required for spreading of Bt8741 colony biofilms

Since we found that the spreading phenotype of Bt8741 requires both *nprR* and *nprRB* genes, the participation of the NprRB signaling peptide was directly assessed. The *nprRB* gene encodes for a 43 amino acid (aa) peptide, that is processed to the 23 aa pro-NprRB during secretion (44, 45); the 23 aa pro-NprRB peptide is further processed extracellularly to the mature form, corresponding to an hepta- or octa-peptide (SKPDIVG or SSKPDIVG) (39). Mature NprRB is reimported and binds to NprR (46). For these experiments, we performed the spreading assay using agar plates supplemented with either the functional 7 aa NprRB SKPDIVG at different concentrations, the 23 aa pro-NprRB peptide or the inactive 5 aa variant SKPDI.

After 7 days, we found that the spreading phenotype of Bt8741 Wt strain was not affected by addition of the synthetic peptides. In contrast, spreading of the Δ*nprR-nprRB*(pMAD-NprR) complemented mutant was restored by the addition of the 7 aa peptide SKPDIVG. This peptide allowed the recovery of spreading at 500 nM, but not at 10 nM, 50 nM or 1 μM. Notably, addition of 500 nM SKPDIVG resulted in a morphology similar to that of the Wt strain at day 7 (Figure 3). Similarly, addition of 500 nM of the 23 aa pro-NprRB rescued spreading in the Δ*nprR-nprRB*(pMAD-NprR) complemented mutant, indicating that this synthetic peptide was processed extracellularly to the mature form and imported into the cells (Figure 3). We did not find recovery of spreading in the Δ*nprR-nprRB*(pMAD-NprR) complemented mutant by the addition of 500 nM of the 5 aa SKPDI; it has been previously shown that this 5 aa peptide does not bind to the NprR receptor *in vitro* (45). Notably, none of these peptides (the mature 7 aa NprRB, 23 aa pro-NprRB or the inactive 5 aa variant) had any effect on spreading of the Δ*nprR-nprRB* mutant. This indicates that the NprR receptor is required to mediate signaling by the 7 aa active NprRB synthetic peptide to achieve this function. Together, these results show that spreading of Bt8741 is regulated by QS, requiring specific signaling by the mature form of the NprRB peptide, and sensing by the NprR receptor.

**Figure 3.**
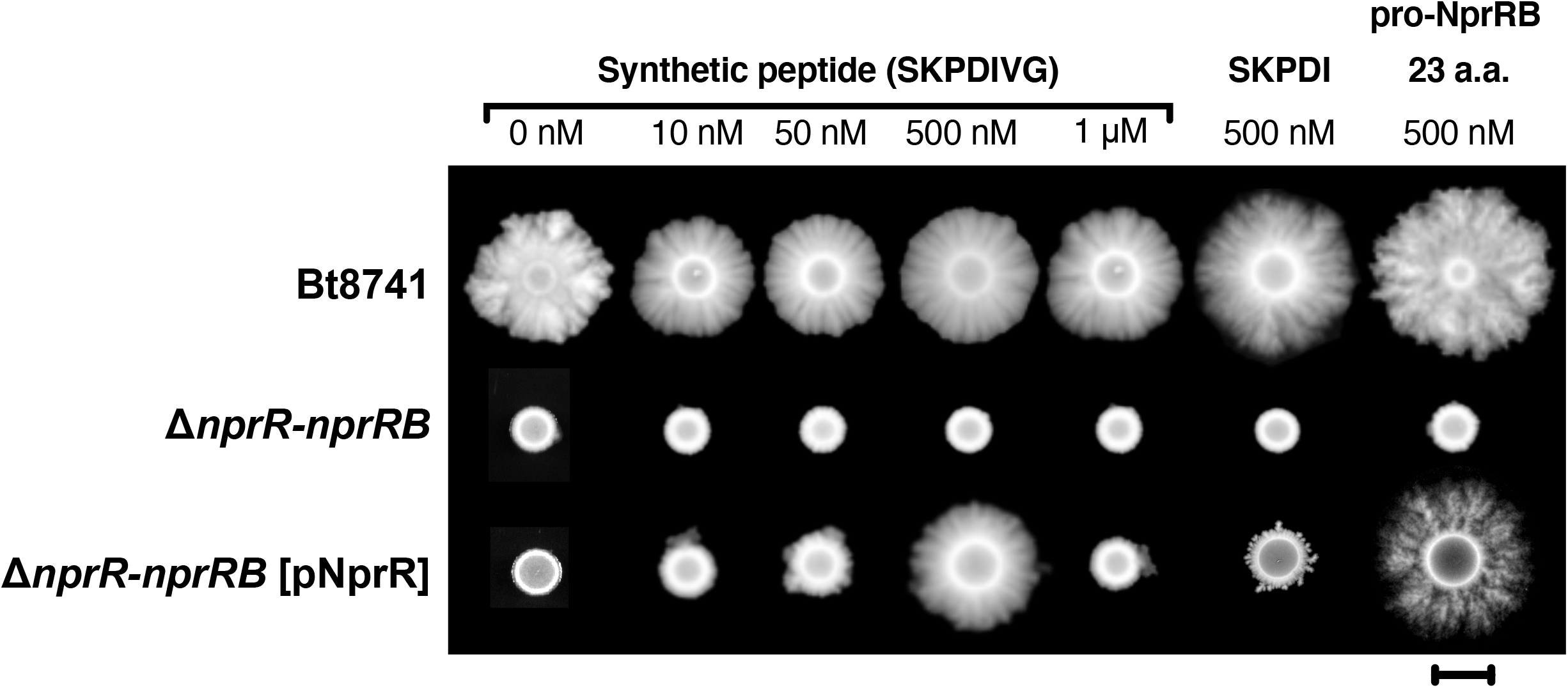
Spreading of Bt8741 in agar plates depends on extracellular signaling by mature NprRB peptide. Colonies of Bt8741 Wt, Δ*nprR-nprRB* mutant and Δ*nprR-nprRB* (pMAD-NprR) complemented mutant were spotted on agar plates containing different concentrations of the SKPDIVG synthetic peptide (10 nM, 50 nM, 500 nM and 1 μM), 500 nM of the inactive form SKPDI or 500 nM of the 23 aa pro-NprRB. All the strains were grown by 7 days before imaging. The scale bar indicates 5 mm. Triplicate for all treatments were performed, and representative images are shown.

### 2.4 Spreading is associated with increased fitness in colony biofilms of Bt8741

Since spreading colonies of Bt8741 Wt gain access to space and probably nutrients that could be unavailable to non-spreading Δ*nprR-nprRB* mutant, we suggested that the spreading phenotype may be associated to an increased growth in colony biofilms. To evaluate growth, the spreading assay was performed, first in colonies of singly-inoculated strains, using Bt8741 Wt, Δ*nprR-nprRB* mutant, Δ*nprR-nprRB* (pMAD-NprR) and Δ*nprR-nprRB* (pMAD-NprR-NprRB) complemented mutants. To determine the growth of these strains, complete colonies were recovered from the agar surface resuspended in PBS, then, total cells in each colony were calculated for each case (Figure 4a). We found that total cells in colonies of the *ΔnprR-nprRB* mutant strain decreased at day 7 in comparison to the Wt strain. While the Wt strain reached 8.0×10^8^ cells per colony, the mutant strain only reached 2.5×10^8^ CFU per colony (Figure 4a). Similarly, the non-spreading Δ*nprR-nprRB* (pMAD-NprR) complemented mutant had decreased growth in these conditions. Finally, the Δ*nprR-nprRB* (pMAD-NprR-NprRB) complemented mutant, which displays the spreading phenotype, reached 8.8×10^8^ CFU per colony at day 7, a similar number to that of the Wt strain (Figure 4a). These results show that spreading phenotype is associated to increased growth of the bacterial population in this *in vitro* model.

**Figure 4.**
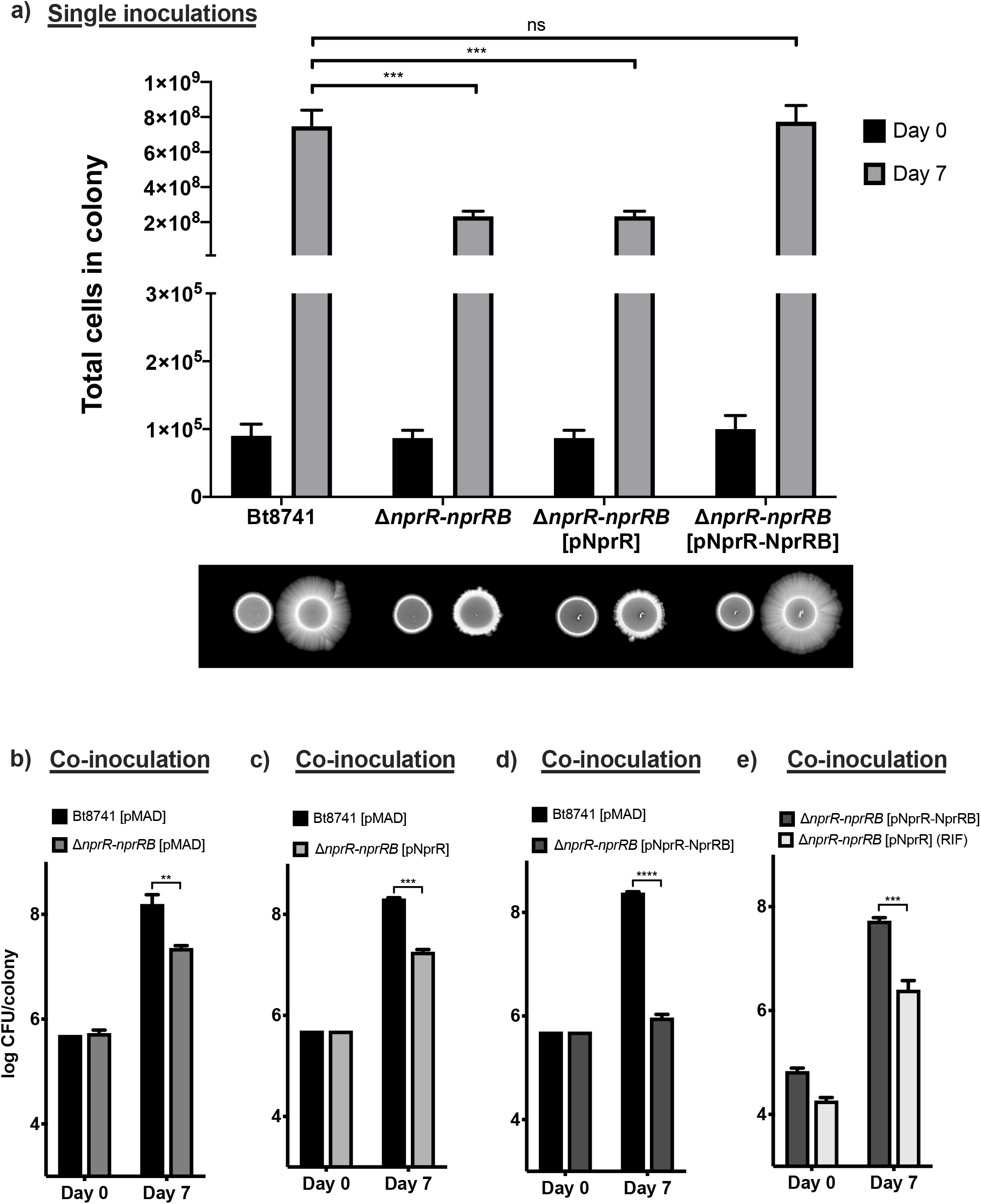
Spreading is associated to increased growth and competitive fitness in colonies of Bt8741. (**a**) Top: Total cells per colony of the Bt8741 Wt, Δ*nprR-nprRB* mutant and Δ*nprR-nprRB* (pMAD-NprR-NprRB) complemented mutant were calculated from single-inoculated colonies after 7 days of growth. Bottom: Representative images of the corresponding spotted colonies at day 0 and 7. (**b**) Competitive fitness of the Wt Bt8741, *B. thuringiensis ΔnprR-nprRB, B. thuringiensis ΔnprR-nprRB* (pMAD-NprR) and *B. thuringiensis ΔnprR-nprRB* (pMAD-NprR-NprRB) strains. Co-inoculated colonies were grown for 7 days; then, total cells of each strain in the colony were calculated. Columns represent the average from 3 replicates ± standard deviation (SD). *, *P* 0.01; ***, *P* < 0.001; ****, *P* < 0.0001; ns, non-significant.

Next, we tested if spreading and growth of Bt8741 was associated to a better competitive fitness, which could explain the fixation of *nprR-nprRB* genes –which control spreading –in natural populations. For this, we performed the spreading assay using 1:1 mixed colony of Wt vs mutant, or Wt vs complemented strains. After 7 days of incubation of the mixed Wt vs mutant colony, the Wt strain reached 2.4×10^8^ CFU per colony, compared to 2.9×10^7^ CFU of *ΔnprR-nprRB* mutant strain (Figure 4b). The result was similar when the Wt strain was co-inoculated with the Δ*nprR-nprRB*(pMAD-NprR) complemented mutant, which does not spread (Figure 4c). When we co-inoculated the Wt strain vs the Δ*nprR-nprRB* (pMAD-NprR-NprRB) complemented mutant, both of which display the spreading phenotype, we found that fitness was not restored in the complemented strain (Figure 4d). We hypothesized that lack of complementation of competitive fitness could be associated to the genetic manipulation in the complemented mutant, which carries a higher copy-number of the *nprR-nprRB* operon. For this reason, we also performed a mixed inoculation using Δ*nprR-nprRB*(pMAD-NprR-NprRB) and Δ*nprR-nprRB*(pMAD-NprR) (RIF) complemented mutants. In this case, the mutant complemented with both NprR-NprRB (which displays the spreading phenotype) had higher fitness than the mutant complemented with only NprR (which does not spread) (Figure 4e). These results suggest that the spreading phenotype controlled by NprR-NprRB, could be related to increased fitness of Bt8741 in natural populations.

### 2.5 Exogenous addition of a surfactant lipopeptide from Bacillus rescues spreading in Bt8741 ΔnprR-nprRB mutant

The spreading phenotype has been reported previously in other *Bacillus* species. Although the mechanisms involved have not been described, it has been proposed that surfactants may be involved in this passive motility (47, 48). In order to evaluate the participation of extracellular compounds in spreading, we performed experiments to assess complementation *in-trans*. First, we tested separate spot-inoculations at different distances (1 to 7 cm) of Bt8741 Wt and Δ*nprR-nprRB* mutant. We expected that diffusion of extracellular metabolites, which are absent in the mutant but produced by the Wt strain, could allow spreading of the mutant strain at closer distances. However, we observed that spreading of the mutant strain was not rescued when it was inoculated adjacent to the Wt strain; accordingly, no differences were observed in the borders of the respective colonies (Supplemental Figure S1). Next, we tested trans-complementation in mixed colonies. For this, we generated GFP-tagged strains for co-inoculations. However, co-inoculation of the Wt strain and the reporter mutant strain Δ*nprR-nprRB* [pHT315 P_*spac*_ ‘*gfp* did not result in trans-complementation of spreading of the mutant strain, which remained localized in the center of the colony (Supplemental Figure S2). In contrast, the GFP-tagged Wt strain spread normally in co-inoculation (Supplemental Figure S2).

Kurstakin gene expression is regulated by the QS NprR-NprRB system (20); hence, we hypothesized that the mutant strain *ΔnprR-nprRB* was defective in the production of this lipopeptide, which could be related to an increased surface tension around the mutant colonies, restricting the spreading phenotype of this strain. To test this, we performed the spreading assay in plates supplemented with the lipopeptide surfactin on the surface of the agar. Surfactin is a lipopeptide from *B. subtilis* (49) with structural and physicochemical similarities to Kurstakin. When surfactin was added, spreading patterns were observed in the mutant strain *ΔnprR-nprRB* after 7 days of incubation (Figure 5a), which were accompanied by cell projections in the edge of the colonies (Figure 5b). This result indicates that the spreading phenotype is related to the production of an extracellular surfactant by Bt8741, probably kurstakin. Further experiments are needed to assess lack of trans-complementation from the Wt strain to the mutant (Supplemental Figures S1 and S2).

**Figure 5.**
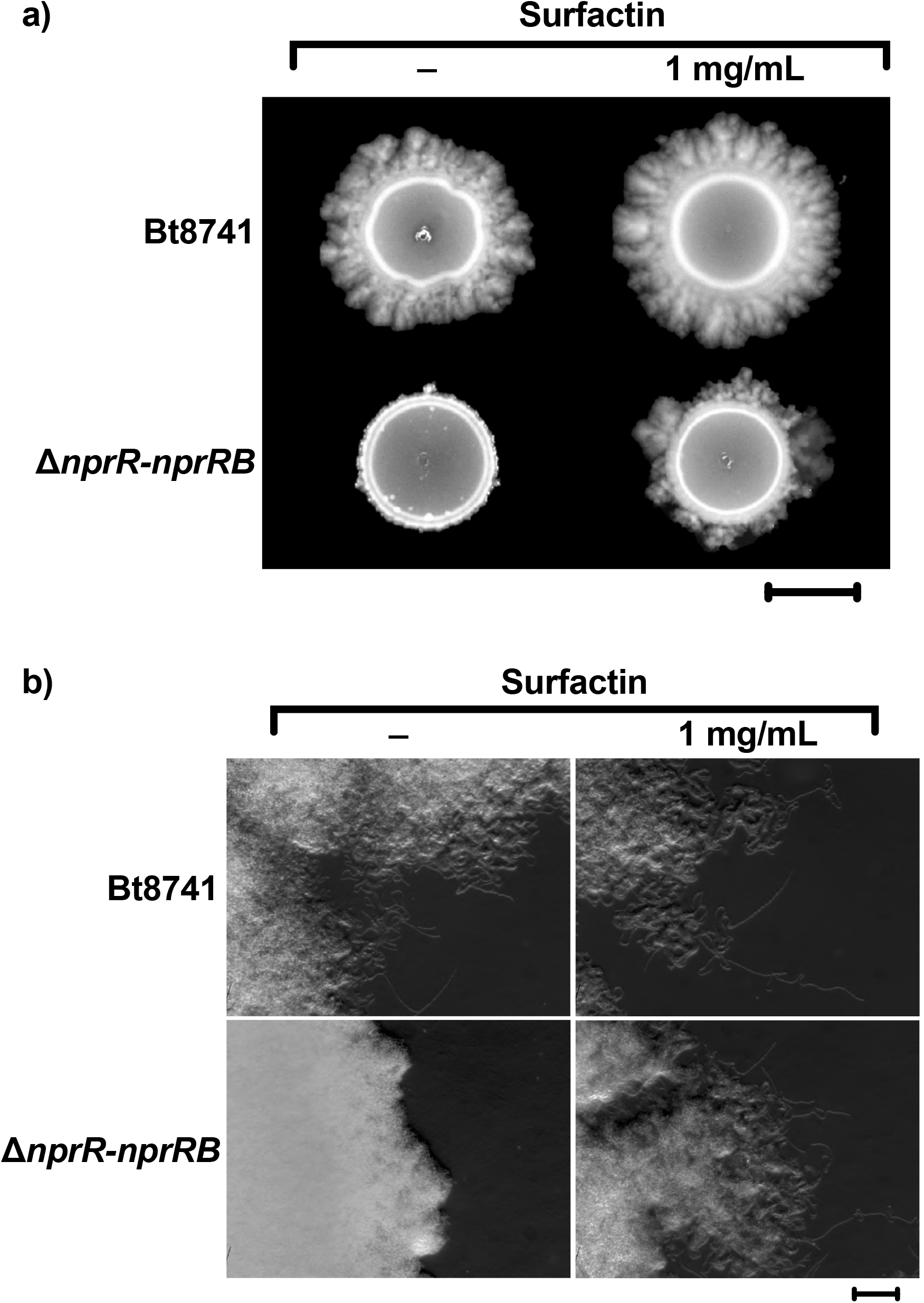
Effect of a surfactin from *Bacillus subtilis* on the spreading phenotype of Bt8741 Wt and *ΔnprR-nprRB* mutant. (**a**) Addition of 1 mg/mL of exogen surfactin restores spreading in the *ΔnprR-nprRB* mutant strain. Scale bar indicates 5 mm. (**b**) Differences in colonies borders and morphology were observed at light field stereo microscope. Scale bar indicates 0.2 mm.

### 2.6 Is the spreading phenotype relevant in nature?

The soil is the main environmental reservoir of species the genus *Bacillus*, where spores can germinate and cells can subsequently grow as saprophytes (50). In order to explore the relevance of spreading in this environment, we assessed the ability of Bt8741 to grow and spread on soil extract. The strains Bt8741 Wt, Δ*nprR-nprRB* mutant, Δ*nprR-nprRB* (pMAD-NprR) and Δ*nprR-nprRB* (pMAD-NprR-NprRB) complemented mutants were spotted individually in a soil extract agar media (see materials and methods). After 7 days, we observed that Bt8741 Wt and Δ*nprR-nprRB* (pMAD-NprR-NprRB) complemented mutant strain exhibited spreading; in contrast, spreading was absent in the Δ*nprR-nprRB* mutant and the Δ*nprR-nprRB* (pMAD-NprR) (Figure 6). This result suggests that *nprR-nprRB* could control the spreading phenotype even in the sole presence of soil nutrients.

**Figure 6.**
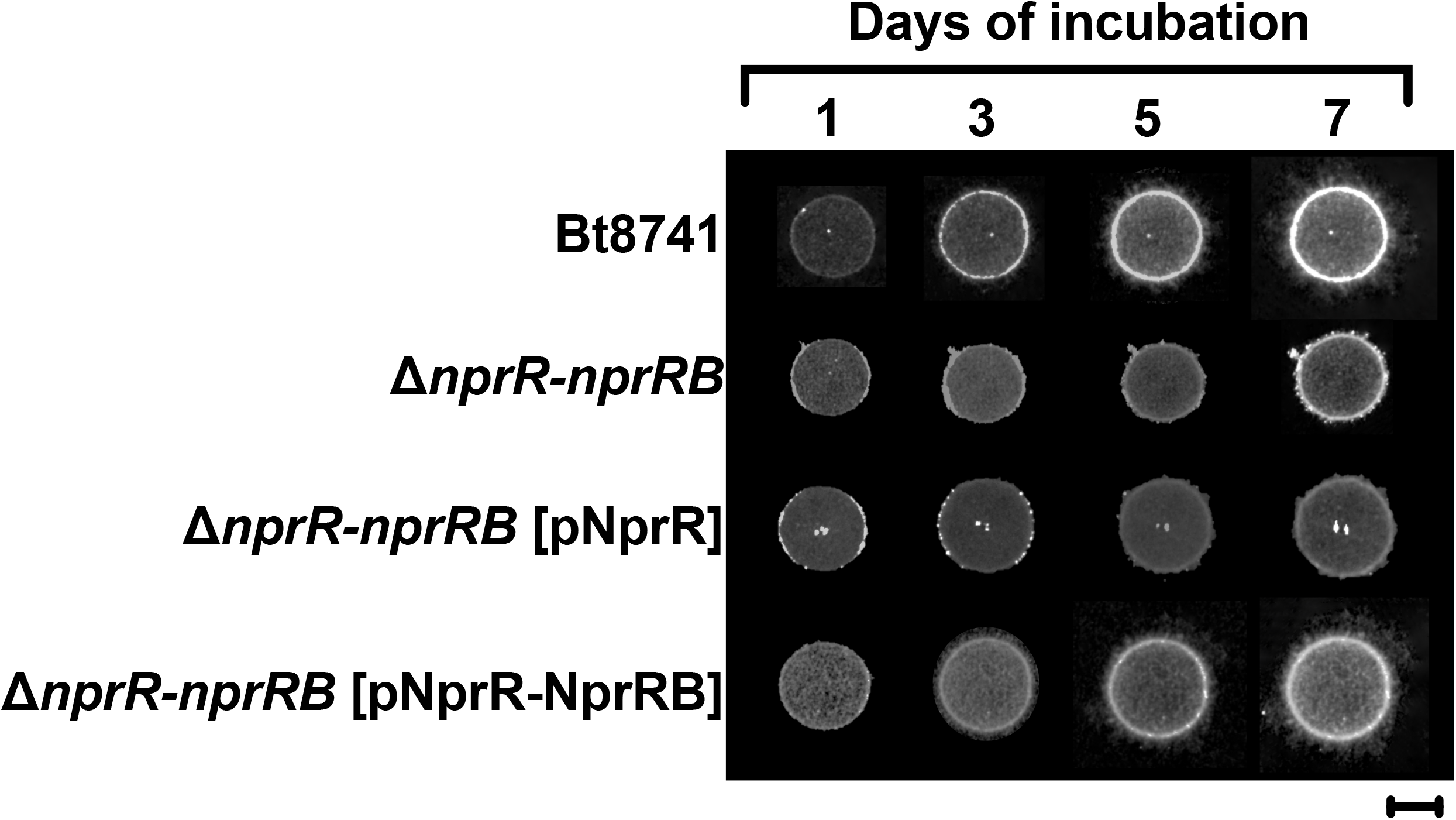
Wt Bt8741 colonies spread in soil nutrients. The strains Wt Bt8741, *B. thuringiensis ΔnprR-nprRB* mutant, *B. thuringiensis ΔnprR-nprRB* (pMAD-NprR) and *B. thuringiensis* Δ*nprR-nprRB* (pMAD-NprR-NprRB) were spotted individually in a soil extract agar media. The scale bar indicates 5 mm.

Bacteria from the genus *Bacillus* are widely distributed in nature among different environments and display an extensive heterogeneity of lifestyles (51). Some studies have assessed the phenotypical diversity associated to the diverse lifestyles of the *Bacillus cereus* group species (16, 52–54), but the spreading phenotype is largely understudied (55–58). To explore if spreading phenotype is a common trait across strains of *B. thuringiensis* and other species of *Bacillus*, the strains Bt8741, BtHD1, BtHD73, *Bacillus cereus* B949 and *Bacillus subtilis* B1015 were spotted individually on different agar media (NA, LB, BHI and TSA) (Figure 7). After 7 days of incubation, a consistent spreading pattern were observed with different magnitudes in all strains and in all media. Since spreading is present in 1) Bt8741 colonies growing in soil nutrients, and 2) a diverse set of *Bacillus* strains from soil, we suggest that this phenotype could be relevant in nature for the survival of the bacteria in soil.

**Figure 7.**
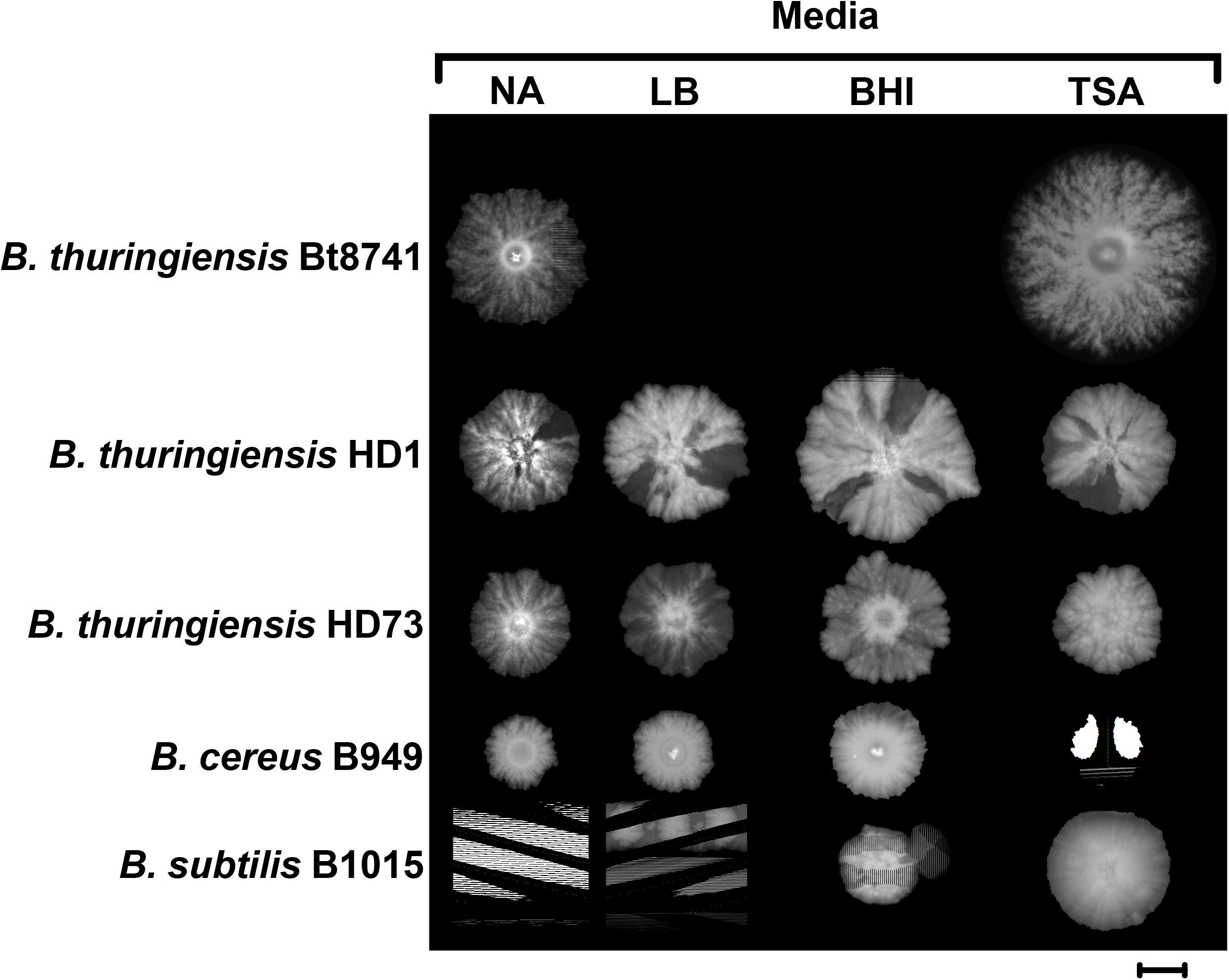
Spreading is a common trait among *Bacillus* species. Preinoculum of the *Bacillus thuringiensis* Bt8741, *Bacillus thuringiensis* HD1, *Bacillus thuringiensis* HD73, *Bacillus cereus* B949 and *Bacillus subtilis* B1015 strains were spotted individually in different agar media to evaluate the spreading phenotype. The scale bar indicates 5 mm.

## 3. Discussion

We investigated the role of the QS system NprR-NprRB on spreading of *B. thuringiensis* Bt8741 and assessed the ecological and evolutionary relevance of this collective trait. Spreading is a passive motility phenotype that has been previously reported in other *Bacillus* (55, 59–61), but the mechanisms underlying this behavior and its regulation are largely unknown. Here we show that spreading in Bt8741 depends on the NprR QS receptor and on signaling by the NprRB mature peptide. We also demonstrate that spreading is associated with increased fitness in colony biofilms of Bt8741, and therefore, the NprR-NprRB QS system is also important for fitness in these conditions. Our results suggest that spreading is mediated by a lipopeptide with surfactant properties, and its synthesis is regulated by NprR-NprRB. Finally, we show that the spreading phenotype occurs in soil nutrients and is conserved in several species of soil *Bacillus*, and hence, this phenotype may be relevant in this niche.

Bacteria use different motility strategies to colonize diverse ecological niches, acquire nutrients and reach favorable environments (62). Active dispersal strategies depend on the presence of flagella, while passive dispersal is flagellum-independent (63–65). One example of a passive dispersal strategy is sliding, which depends on the secretion of surfactants that reduce surface tension and facilitate the bacterial translocation (47, 66, 67). Bacteria from the genus *Serratia, Pseudomonas, Legionella, Sinorhizobium, Salmonella, Mycobacterium* and *Bacillus* spread by the sliding strategy, which is allowed by the secretion of lipopeptides, glycopeptidolipids and exopolysaccharides (55, 68). This collective behavior allows access to nutrient sources that could not be reached by single individuals (69). For example, some bacterial species colonize and spread into intracellular and extracellular plant tissues as endophytes, reaching the roots, stem, flowers, leaves, fruit or seeds; all these tissues are abundant in carbohydrates, amino acids and other nutrients required for optimal bacterial development (70, 71).

Despite the extensive research about *B. thuringiensis*, most of the work has focused on the relevance of *B. thuringiensis* for pest control in agriculture. Its ecology related with other lifestyles (free or plant-associated) and the interaction with other microorganisms or plant hosts have been poorly addressed, since this species has mainly considered an entomopathogenic or saprophytic bacterium (10, 31, 32, 54, 72, 73). In turn, the mechanisms that *B. thuringiensis* uses to spread in soil or other habitats are still unknown. It has been reported that it occurs mainly passively by the action of rain, wind, water, plant growth, and by the presence of spores in insect cuticles and feces (74–81). However, *B. thuringiensis* can also colonize diverse tissues of plants, being found mainly in the roots where exudates rich in carbohydrates and amino acids are secreted and favor its growth (82). The spreading phenotype may be relevant during colonization of these niches, increasing survival after translocation across biotic or abiotic surfaces, allowing access to nutrients and escape from unfavorable environments (83–85).

In a recent work, we reported that four Rap proteins (RapC, RapF1, RapF2, and RapLike) negatively control the spreading in *B. thuringiensis* (28). In contrast, here we show that NprR favors dispersion of colony biofilms. Redundant control of spreading by several QS receptors, point to the fact that this phenotype is relevant in nature. Additionally, the regulation of spreading by several QS regulators could be even more complex in nature, where other diverse signals, cues and stimuli may be found.

Previous studies have shown that the NprR QS receptor of *B. thuringiensis* activates the transcription of genes related with the saprophytic and necrotrophic lifestyle and the sporulation initiation through modulation of the Spo0A-phosphorelay by protein-protein interaction with Spo0F (20, 40, 43). However, the evidence accumulated indicates that different functions of NprR are evident in diverse conditions or ecologic context. On one hand, experiments performed in liquid media showed no growth differences between Bt8741 Wt and the *ΔnprR-nprRB* mutant, while only a slight temporal delay on sporulation was found (40). In contrast, in experiments performed in an insect host model, NprR is essential to assure the growth and survival of *B. thuringiensis* through the activation of genes related to necrotrophism, also contributing to sporulation after the necrotrophic phase (20, 86). Here, using a colony biofilm model, we find that a *nprR* deletion has no effect on sporulation and a moderate effect on growth fitness. Moreover, the most prominent effect of *nprR* deletion in this context appears to be related with the production of a surfactant molecule, which allows spreading by reducing surface tension and let effortless movements (87). Surfactants also contribute to biofilm formation and increased microbial competition in bacteria from the *B. cereus* group (88), suggesting that NprR-NprRB QS systems may be relevant in other, not yet addressed conditions. In this sense, our work offers new insights into the control of functions that are relevant for *B. thuringiensis* lifestyle in nature.

The NprR regulon includes many degradative enzymes and proteins involved in the synthesis of kurstakin, a lipopeptide with antifungal and biosurfactant properties (20, 89). The synthesis of kurstakin in *B. thuringiensis* may also be related to the facilitation of spreading of vegetative cells to reach new nutrient sources and guarantee the survival of *B. thuringiensis* cells in different environments. In *B. thuringiesis* Bt407, four genes (*krsEABC*) are required for the synthesis of kurstakin (20, 90). Here we show that surfactin mimics the function of a *B. thuringiensis-produced* lipopeptide, which appears to mediate spreading. The lipopeptide surfactin shares similar structure and properties with lipopeptide kurstakin, and both act as surface tension reducers (87). Additionally, it has been shown that the lipopeptide surfactin acts as a signal molecule in *B. subtilis*. Surfactin promotes phosphorylation of Spo0A via the membrane protein histidine kinase KinC, in turn activating the SinI-SinR pathway that leads to biofilm matrix production (91). Together, evidence suggest that spreading of Bt8741 requires the activation of kurstakin synthesis by NprR-NprRB; this molecule may act both as surfactant and as a signal molecule for promoting the production of other extracellular compounds via sensor kinases and the Spo0A-phosphorelay. This is consistent to our observation that the sole exogenous addition of surfactin is sufficient for restoring spreading in the Δ*nprR-nprRB* mutant strain. However, extracellular compounds produced by Wt Bt8741 were not sufficient for restoring spreading of Δ*nprR-nprRB* mutant *in-trans*. This phenomenon could be associated to a kin-recognition mechanism between the public-good-producing and non-producing population (92, 93), which evolve to prevent exploitation by the non-producers.

## 4. Conclusions

Studies on *Bacillus thuringiensis* have focused mainly on its biotechnological applications. Only recently, the adoption of *B. thuringiensis* (and other species from the *B. cereus* group) as model for studying highly complex QS-control of cell processes, has resulted in the generation of knowledge on its ecology, albeit mostly as an insect pathogen. Bacteria evolve to thrive in changing environments and spreading is a phenomenon that appears to be relevant for adaptation of *B. thuringiensis* to its surroundings. In addition to the functions of NprR-NprRB reported in previous work, the control of spreading highlights the relevance of this QS system for the interaction of *B. thuringiensis* and other *Bacillus cereus* group species with their environment. Further studies should aim at obtaining direct observations of spreading in nature as well as uncovering the molecular mechanisms relevant for *B. thuringiensis* during its free-lifestyle, in root-associated biofilms, or as plant endophytes.

## 5. Materials and Methods

### Bacterial strains, plasmids, synthetic peptides, media and culture conditions

All strains and plasmids used in this study are listed in Table A1. *B. thuringiensis* strain 8741 (Bt8741) was used as wild type strain (39). Mutants in the *nprR-nprRB* locus and complemented strains were described in a previous work (40). Luria-Bertani (LB) broth (5 g L^-1^ NaCl, 5 g L^-1^ yeast extract and 10 g L^-1^ tryptone) and nutrient agar (8 g L^-1^ nutrient broth, 15 g L^-1^ agar) were routinely used at 30 °C for *B. thuringiensis* cultures. Peptide variants were designed from the amino acid sequence of NprRB, synthesized by GeneScript®, and added to cultures at variable concentrations. When needed, erythromycin (5 μg mL^-1^), spectinomycin (250 μg mL^-1^), rifampicin (50 μg mL^-1^) or surfactin (1 μg μL^-1^) (Sigma-Aldrich, St. Louis, MO, USA) was added.

### Spreading assay in agar plates

An agar plate assay was designed to follow the motility phenotype of *B. thuringiensis* strains. Colonies of *B. thuringiensis* Wt, mutant or complemented strains (Table 1) were picked into 10 mL of LB medium with erythromycin and grown overnight at 200 rpm. Then, 5 μL of these preinoculum cultures, containing ~10^6^ cells ml^-1^ were spotted on a diluted nutrient broth, 0.8 g L^-1^ of media were utilized instead of 8 g L^-1^. Plates were incubated at 30 °C for 7 days and imaged daily using a gel documentation system (Gel Doc^™^ XR+, BIORAD). When needed, the diametral growth of each colony was measured using the Image Lab™ Software (BIORAD) and correlated with the mobility capacity based on its genotype. All experiments were performed in three replicates per strain.

### Growth fitness assay

To assess growth fitness, we quantified total CFU in droplets inoculated at Day 0 and the final CFU counts at day 7 in the biofilm assay. For measuring initial bacterial cells in colonies, 10-fold serial dilutions of the preinoculum were plated, incubated overnight at 30°C; resulting colonies were counted and the total CFUs in the inoculated droplet were calculated. For measuring final bacterial cells in the developed biofilm, the entire colony was cut off the agar surface, and cells were suspended in 2 ml of phosphate buffered saline (PBS; 1.7mM KH2PO4, 5mM Na2HPO4, 150mM NaCl) by intense vortexing for 1 min. The cell suspension was checked for agglomerates by microscopy, and then 10-fold dilutions of this suspension were plated. After overnight incubation, colonies were counted and total cells in the biofilm at 7 days were calculated. All experiments were performed in triplicates.

### Mixed biofilm experiments

Co-culture experiments were performed to test if the motility phenotype could be complemented extracellularly *in trans*. For this, overnight cultures were washed and combined in pairs (Wt & mutant strain, Table 1) in equal density (10^6^ cells mL^-1^). Then, 5 μL of each mixture was spotted on diluted nutrient agar and incubated for 7 days until biofilm was developed. Relative abundance of each strain in the mixed biofilm was measured as follows: samples of each mixed biofilm were picked with a wooden stick from three sites of the colony (center, edge and the midpoint). Each sample was suspended 100 μL of PBS, and then 10-fold dilutions of each were plated in nutrient agar with erythromycin, and in nutrient agar with spectinomycin and erythromycin. After 18 hours of incubation at 30°C, colonies were counted. SpecR colonies correspond to mutant strains, and Wt CFU were obtained from subtracting total CFU - SpecR CFU.

In order to follow growth of the mutant strain in the mixed culture, the *ΔnprR-nprRB* mutant was transformed with pHT315-Pspac-*gfp*, obtaining a constitutive GFP fluorescent strain. This strain was mixed with the Wt strain and fluorescence of the mixed biofilm was followed by imaging with a ChemiDoc XRS+ (BIORAD, CA, USA) using transmitted light and 62 mm standard emission filter. Images were overlapped in image J.

### GFP-labeling of *B. thuringiensis* Bt8741

For the generation of fluorescent *B. thuringiensis* strains, the *gfp* gene was amplified from plasmid pMUTIN-GFP (41), using primers DS13-Fwd (5’-TCT TCT AGA TTG ACT TTA TCT ACA AGG TGT GGC ATA ATG TGT GTA ATT GTG AGC GGA TAA CAA TTA AGG AAG GAG ATA TAC ATA TGG C-3’) and DS2-Rev (5’-TCT TCT AGA CAA TCA CGA AAC AAT AAT TG-3’); where DS13 contains an upstream sequence corresponding to the constitutive promoter Pspac. The PCR product was inserted into the *XbaI* restriction site of the plasmid pHT315 (42). Then, this plasmid was used for transformation of electrocompetent Bt8741 Wt and *ΔnprR-nprRB* mutant strain, as reported previously (40).

### Microscopy

An Axio Zoom.V16 stereo microscope with an Axiocam 105 color incorporated (Carl Zeiss Microscopy, Oberkochen, Germany) was used to obtain images from the edge of spreading colony biofilms. Samples of the strains (Bt8741 Wt, Δ*nprR-nprRB* mutant and complemented strains) were inoculated as previously described. Images of 180X resolution were obtained to visualize the border of each colony. Each strain was analyzed for triplicated to demonstrate the reproducibility of the colony patern.

## Author Contributions

Conceptualization, J.R. and A. V-F.; methodology, A. V-F. and J.R.; software, A. V-F. and J.R.; validation, A. V-F. and J.R.; formal analysis, A. V-F. and J.R.; investigation, A. V-F., M.T. and J.R.; resources, J.R. and M.T.; data curation, A. V-F. and J.R.; writing—original draft preparation, A. V-F.; writing—review and editing, A. V-F., M.T. and J.R.; visualization, A. V-F., M.T. and J.R.; supervision, J.R.; project administration, J.R. and M.T.; funding acquisition, J.R. and M.T. All authors have read and agreed to the published version of the manuscript.

## Acknowledgments

This work was supported by CONACYT (Mexico) grant 267837 to M.D.L.T. A fellowship from CONACYT (No. 449242) was given to A.V-F. Daniel Perez constructed the plasmid for GFP labeling of Bt. Jose Angel Huerta provided technical assistance in macroscopic fluorescence imaging.

