## Supplementary Material for "The quorum sensing system NprR-NprRB contributes to spreading and fitness in colony biofilms of *Bacillus thuringiensis*"

**Supplementary Figures:**

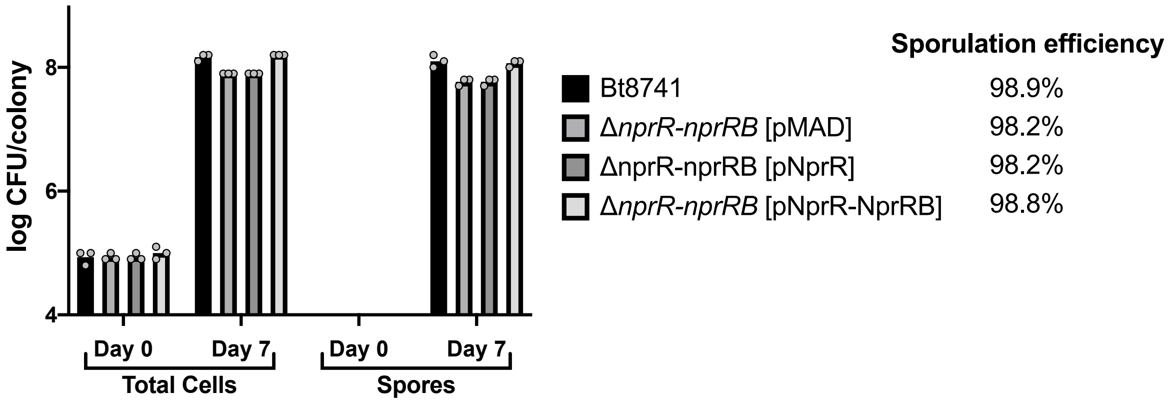


**Supplemental Figure S1. Sporulation efficiency in Wt, mutant and complemented strains.** Bt8741 WT strain, *B. thuringiensis* Δ*nprR-nprRB* mutant, *B. thuringiensis* Δ*nprR-nprRB* (pMAD-NprR) and *B. thuringiensis* Δ*nprR-nprRB* (pMAD-NprR-NprRB) complemented strains were picked in 5 mL of LB media and grow overnight in orbital shaker at 30 ºC and 200 rpm. When needed, erythromycin (5 µg $mL^{-1}$) was added. Cell of each culture was washed and resuspend in phosphate buffered saline (PBS; 1.7mM KH2PO4, 5mM Na2HPO4, 150mM NaCl). Then, 5 µL of each strain containing ~$10^6$ cells $ml^{-1}$ were spotted in the center of a diluted nutrient broth agar media, (0.8 g $L^-$ of and 15 g $L^-$ of agar agar), plates were incubated at 30 ºC for 7 days. After incubation, the colony of each strain was cut off the agar surface, cells were resuspended in 5 ml of phosphate buffered saline by vortex. 10-fold ($10^{-1}$-$10^{-6}$) serial dilutions of each resuspension were plated and incubated overnight at 30ºC to calculate total CFU/colony. To determinate the number of spores, 1 mL of each resuspension were heated for 15 min at 80 ºC to remove non-sporulated forms and obtain thermoresistant cells. Then, serial dilutions were plated as described above. CFU/colony were calculated and to estimate sporulation efficiency, CFU/colony of thermoresistant cells were subtracted from the total CFU/colony. Results are expressed as percentages. All the experiments were repeated three times and are represented in columns as dots.

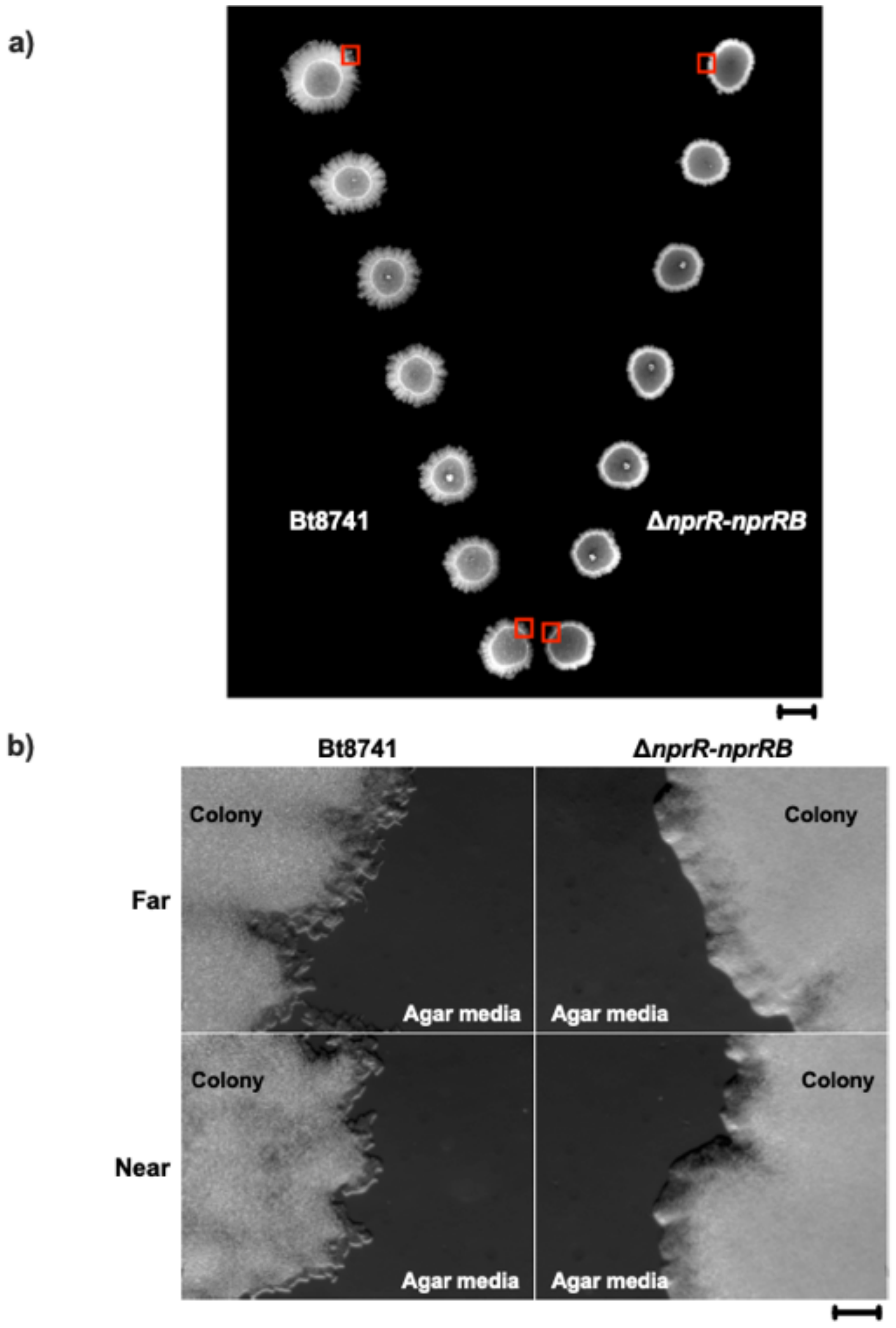


**Supplemental Figure S2. Proximity assays for *in-trans* complementation.** Preinoculum of the Bt8741 WT strain and *B. thuringiensis* Δ*nprR-nprRB* mutant were grown overnight at 30 ºC and 200 rpm in an orbital shaker. Then 5 µL of each strain was spotted in diluted nutrient broth agar media, (0.8 g $L^{-}$ of and 15 g $L^{-}$ of agar agar) separated from a distance of 1 to 7 cm at a concentration of ~$10^{6}$ cells $ml^{-1}$. Plates were incubated at 30 ºC for 7 days. Subsequently, colonies were imaged using a gel documentation system (Gel Doc™ XR+,

BIORAD). Border colonies were imaged using an Axio Zoom.V16 stereo microscope with an Axiocam 105 color incorporated (Carl Zeiss Microscopy). The scale bar in supplemental figure S2a indicates 5 mm. The scale bar in supplemental figure S2b indicates 0.2 mm.

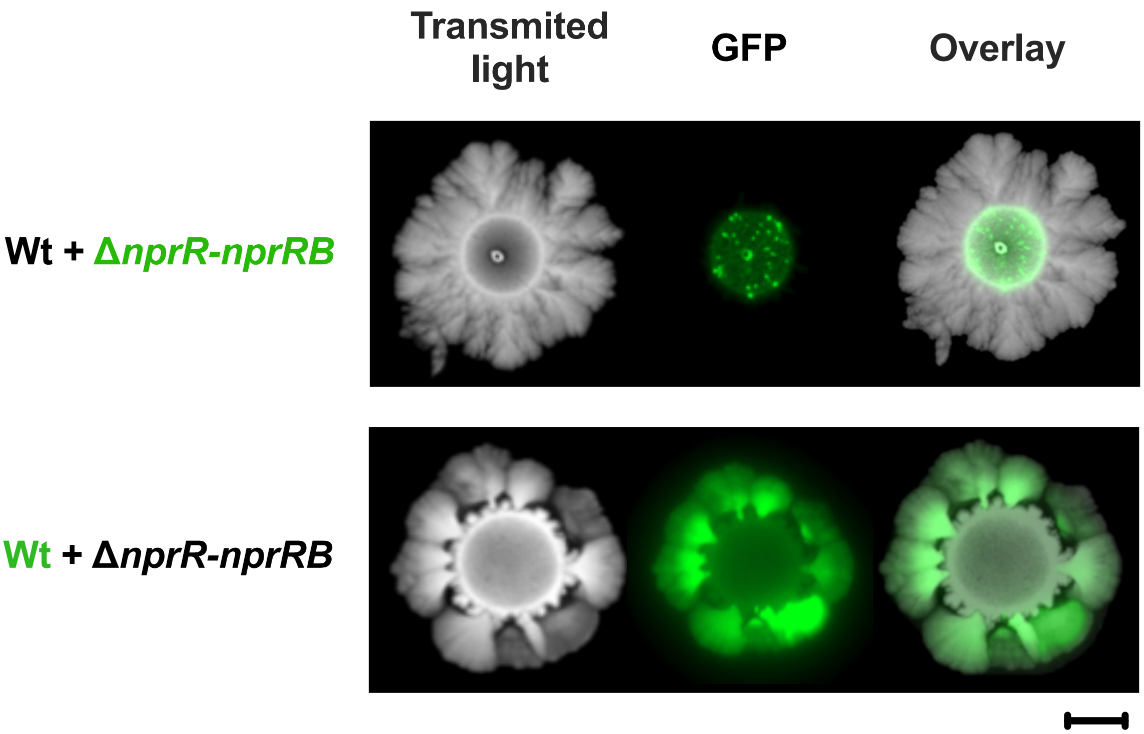


**Supplemental Figure S3.** Participation of extracellular compounds in strain co-inoculations. Preinoculum of the Bt8741 Wt, *B. thuringiensis* Δ*nprR-nprRB* mutant, Bt8741 [pHT315 Pspac'gfp] and *B. thuringiensis* Δ*nprR-nprRB* mutant [pHT315 Pspac'gfp] were incubated overnight at 30 ºC in an orbital shaker at 200 rpm. Co-inoculations of Bt8741 Wt + *B. thuringiensis* Δ*nprR-nprRB* [pHT315 Pspac'gfp] and Bt8741 [pHT315 Pspac'gfp] + Δ*nprR-nprRB* mutant were performed to evaluate the possibility of complementation in trans. Co-inoculation colonies were imaged using a ChemiDoc XRS+ using transmitted light and 62 mm standard emission filter. The scale bar indicates 5 mm.

**Supplementary Table:**

**Supplemental Table S1. Strains and plasmids used in this study.**

| Strain | Description | Reference |
|---|---|---|
| *E. coli* TOP10 | | |
| *E. coli* MC1061 | | |
| *B. thuringiensis* Bt8741 | Wild type strain | (39) |
| *B. thuringiensis* Δ*nprR-nprRB* | Mutant strain, genetic exchange Δ*nprR-nprRB::specR* | (40) |
| *B. thuringiensis* Δ*nprR-nprRB* [pMAD-NprR] | Mutant strain, SpecR, EriR, complemented with NprR | (40) |
| *B. thuringiensis* Δ*nprR-nprRB* [pMAD-NprR-NprRB] | Mutant strain, SpecR, EriR, complemented with NprR and NprRB | (40) |
| *B. thuringiensis* Bt8741 [pHT315 Pspac'gfp] | Wild type strain, EriR, constitutive GFP | This work |
| *B. thuringiensis* Δ*nprR-nprRB* [pHT315 $P_{spac}$'*gfp*] | Mutant strain, genetic exchange Δ*nprR-nprRB::specR*, EriR, GFP transcriptional fusion | This work |
| | Plasmids | |
| pHT315 $P_{spac}$'*gfp* | Plasmid for carrying GFP transcriptional fusions | This work |
| pMUTIN (GFP) | Plasmid for GFP amplification sequence | (41) |
| pMAD | Plasmid for homologous recombination and complementation | (94) |
